# Soluble NS1 antagonizes IgG- and IgA-mediated monocytic phagocytosis of DENV infected cells

**DOI:** 10.1101/2022.12.17.520876

**Authors:** Mitchell J. Waldran, Adam D. Wegman, Lauren E. Bahr, Jeffrey R. Currier, Adam T. Waickman

## Abstract

Dengue virus (DENV) is endemic in over 100 countries, infecting an estimated 400 million individuals every year. Infection with DENV raises a significant antibody response, primarily consisting of antibodies targeting viral (structural) proteins. However, not all DENV antigens are part of the virion itself, as the DENV genome encodes several non-structural (NS) proteins. One of these, NS1, has been shown to be antigenic and is expressed on the membrane of DENV-infected cells. IgG and IgA isotype antibodies that bind NS1 are detectable in serum following DENV infection and are also capable of interacting with Fc receptors expressed on professional phagocytes. Our study aims to determine if NS1-binding IgG and IgA isotype antibodies contribute to the clearance of DENV-infected cells by professional phagocytes through antibody mediated phagocytosis/trogocytosis. Using an in vitro model of trogocytosis we observed that both IgG and IgA isotype antibodies can facilitate facilitating monocytic uptake of DENV NS1 expressing plasma membrane in an additive fashion. This process was dependent on the expression of FcγRI (CD64) and FcαR (CD89) for IgG and IgA mediated membrane uptake, respectively. Furthermore, this process was antagonized by the presence of soluble NS1, suggesting that the production of soluble NS1 by infected cells may serve as an immunological chaff, thereby antagonizing opsonization and clearance of infected cells by NS1-specific IgG and IgA isotype antibodies.

## Introduction

Dengue virus (DENV) infects an estimated 400 million individuals annually, resulting in 100 million clinically apparent infections and at least 40,000 deaths (1–3). Consisting of four immunologically and genetically distinct serotypes (DENV-1, -2, -3 and -4), DENV is transmitted by the bite of infected mosquitos belonging to the genus *Aedes* (1, 4, 5). Primary DENV infections are typically mild; causing fever, rashes, nausea, vomiting, and myalgias and arthralgias (1, 5, 6). However, a fraction of heterologous secondary infections progress to severe dengue, characterized by plasma leakage or bleeding leading to shock or respiratory distress with substantial mortality risk in the absence of appropriate supportive care (7–9). The leading mechanistic explanation for the increased disease risk associated with post-primary dengue is a process known antibody dependent enhancement (ADE), wherein DENV-reactive antibodies generated by a primary DENV infection facilitate the infection of cells that are normally not susceptible to the virus via Fcγ receptor-mediated endocytosis (10–12). This process is thought to increase both the number of cells infected by DENV in immunologically primed individuals relative to their naïve counterparts, but also to elicit a more significant acute inflammatory response to infection, leading to increased immunopathology (13). This distinct epidemiological feature of dengue has severely hindered the development of safe and effective dengue vaccines despite over 50 years of concerted effort by the scientific community.

Antibodies directed against the structural components of the DENV virion – namely the E and prM proteins – are responsible for both neutralizing and enhancing the infectivity of the virus depending on the abundance of antibody and the exact antigenic epitope engaged by an antibody (14). However, these are not the only antigenic viral proteins expressed by DENV-infected cells. Non-structural protein 1 (NS1) is a 55kDa glycoprotein that is thought to play a role in several steps of dengue virion assembly. Initially synthesized as a monomer, intracellular NS1 (iNS1) rapidly forms a membrane-associated homodimer in the ER and serves as core component of the viral replication complex (15, 16). Dimeric NS1 can also be found on the membrane of infected cells (mNS1) (17–20), and is secreted by infected cells into the extracellular space in higher-order oligomeric forms (21, 22). Once in circulation, secreted NS1 (sNS1) is thought to be a mechanistic trigger of vascular leakage symptoms in DHF/DSS by binding and destabilizing vascular endothelial cells and by activating the innate immune system through direct engagement of toll-like receptors (23, 24). Accordingly, reducing the immunostimulatory capability of sNS1 is a leading avenue of anti-dengue therapeutic research.

Antibodies capable of binding both sNS1 and mNS1 can be found in circulation shortly after the appearance of antibodies directed against the DENV virion, with IgM, IgA, and IgG all represented (25–27). As with DENV virion-binding antibodies, NS1-binding IgM and IgA appear transiently after infection, while NS1-specific IgG persists in circulation (25). How these different antibody isotypes functionally contribute to the resolution of an acute DENV infection and provide protection from subsequent infection is unclear.

Antibodies of all isotypes are in theory capable of binding and sequestering sNS1 – thereby reducing the immunopathogenic properties of the protein – while IgM and IgG isotype antibodies can facilitate lysis of DENV-infected cells via complement fixation (28). However, IgG and IgA isotype antibodies are additionally capable of interacting with their cognate Fc receptors on innate immune cells, thereby facilitating an array of Fc-receptor dependent effector function (29). This includes antibody dependent cellular cytotoxicity (ADCC) – facilitated by FcγR expressing NK cells – as well as antibody-dependent cellular phagocytosis (ADCP). Of note, multiple professional phagocytes, including monocytes, macrophages, and dendric cells, express both FcγRs and FcαR, meaning they are potentially capable of utilizing both IgG and IgA isotype antibodies for the phagocytic clearance of opsonized viral antigen and virally infected cells (28). However, what role – if any – opsonizing IgG and IgA isotype anti-mNS1 antibodies play in the phagocytic clearance of DENV-infected cells remains unclear.

To fill this knowledge gap, we developed an *in vitro* model of NS1-directed phagocytosis and a pair of idiotope-matched NS1-reactive IgG and IgA isotype mAbs. Using this system, we observed that both IgG and IgA isotype antibodies are capable of facilitating monocytic uptake of DENV NS1 expressing cellular material in an additive fashion. This process was dependent on the expression FcγRI (CD64) and FcαR (CD89) for IgG and IgA mediated membrane uptake, respectively. Furthermore, this process was antagonized by the presence of sNS1, suggesting that the production of sNS1 by infected cells may serve as an immunological smoke screen, limiting opsonization and clearance of infected cells by opsonizing antibodies.

## Materials and Methods

### Cell lines and primary human PBMC isolation

A DENV-2 NS1 expressing cell line was generated by electroporating CEM.NK^R^ cells with a linearized plasmid containing the codon-optimized sequence shown in **Supplemental figure 2**, followed by antibiotic selection, single-cell sorting. A DC-SIGN (CD209) expressing cell line was generated by transfecting CEM.NK^R^ cells with a linearized plasmid containing the codon-optimized sequence of DC-SIGN (Genbank accession = NM_021155.4), followed by the same selection and subcloning process as above. Surface expression of DENV-2 NS1 and DC-SIGN was confirmed by flow cytometry prior to each assay. PBMC were collected and isolated from normal healthy volunteers using BD Vacutainer CPT tubes containing sodium heparin. SUNY Upstate Medical University Institutional Review guidelines for the use of human subjects were followed for all experimental protocols in the study.

### Monoclonal antibodies and serum

The variable regions from the heavy and light chains of the previously described DENV-2 reactive murine monoclonal antibody 1G5.3 (30) were codon optimized, synthesized in vitro, and subcloned into an expression vector containing the human IgG1 or IgA1 Fc region by a commercial partner (Genscript). Transfection grade plasmids were purified by maxiprep and transfected into CHO cells. The cell supernatants collected for antibody purification. Following centrifugation and filtration, the cell culture IgG supernatant was purified using MabSelect SuReTM LX, and the IgA supernatant purified using CaptureSelectTM IgA affinity matrix. Antibody lot purity was confirmed by SDS-PAGE to ≥85% purity, and the final concentration determined by 280 nm absorption. Dengue virus immune serum was obtained from a commercial source (SeraCare, Milford, MA)

### Viruses

DENV-2 (strain S16803) was propagated in Vero cells and titrated on DC-SIGN expressing CEM.NKR cells prior to use.

### NS1 ELISA

96-well ELISA plates were coated with 50uL of a 2ug/mL solution of purified DENV-2 NS1 (Native Antigen, DENV2-NS1-100) or DENV-1 NS1 (Native Antigen, DENV1-NS1-100) diluted in carbonate/bicarbonate buffer and incubated overnight at 4° C. The wells were washed 4 times with PBS-T and blocked with blocking buffer for 30 minutes at RT. Blocking buffer was removed, and monoclonal antibodies were prepared at 10ug/mL top concentration followed by 4-fold serial dilutions. The plates were washed 4 times with PBS-T. Peroxidase-labeled anti-human IgG (Southern Biotech 2044-05) or IgA (Biolegend 411002) were added to the wells and incubated for 1 hour at RT. The plates were washed 4 times with PBS-T, and TMB substrate was added to the wells for 10 minutes, and read on a Biotek µQuant plate reader at 450nm.

### Antibody-Dependent Trogocytosis Assay

CEM.NK^R^ and DENV-2 NS1-expressing CEM.NK^R^ cells were stained with PKH26 at a final concentration of 3.0×10^−6^M resuspended in Diluent C for 5 minutes at room temperature. Cells were then washed twice with PBS + 2% FBS prior to opsonization at 4°C with sera (1:500 dilution) or anti-NS1 monoclonal antibodies (mAbs) (1µg/mL). After opsonization, 2×10^4^ of the labeled CEM.NK^R^ cells in a volume of 100µL were added to a well of a 96 well plate, and mixed with 100uL of purified PBMC at 2×10^5^ cells/mL. The cells were incubated together for 3 hours at 37LC. The cells were then washed twice, stained with anti-human CD14 (Biolegend 301818 clone M5E2), and analyzed on the LSRII flow cytometer (BD Biosciences). Purified soluble DENV-1 NS1 protein (Native Antigen DENV1-NS1-100) and DENV2-NS1 (Native Antigen DENV2-NS1-100) was added as a competitive inhibitor for some experiments at a final concentration of 1µg/mL.

### FcR Blocking Assay

Following the same trogocytosis assay as above, before incubating PBMCs with NKR cells, PBMCs are incubated with either anti-CD89 (Biorad MCA1824, clone MIP8a), -32 (Stemcell Technologies 60012 clone IV.3), or -64 (Biolegend 305002 Clone 10.1).

### Microscopy

PKH26 labeled CEM.NKR cells or sorted PKH26+CD14+ monocytes were purified using an BD FACSAria III Cell Sorter (BD Bioscience). Cells were allowed to settle on poly-L lycine coated plates before being imaging on a Nikon Eclipse TI microscope with a Nikon Digital SIGHT DS-Qi1Mc camera, Nikon Intesilight C-HGFIE mercury lamp. Image collection was performed using NIS-Elements AR 4.60.00 (Nikon), and analyzed using FIJI 2.1.1 (31).

### Quantification of surface Fc receptor abundance

PE-conjugated calibration beads (Bang laboratories) were analyzed by flow cytometry at the same time as live cells stained with saturating concentrations of PE-conjugated antibodies against Fc receptors of interest. Linear regression calculation was performed using the isotype-subtracted Mean Fluorescence Intensity (MFI) values from the PE calibration beads and known number of PE molecules per bead. The resulting equation was used to calculate the number of cytokine receptors present on live cells, using the isotype-subtracted PE MFI values from each Fc receptor analyzed. The anti-FcR antibodies used were: PE anti-human CD16 (Biolegend 302007 clone 3G8), anti-human CD32 (Biolegend 303205 clone FUN-2), PE anti-human CD64 (Biolegend 305007 clone 10.1), PE anti-human CD89 (Biolegend 354103 clone A59), PE mouse IgG1 isotype control (Biolegend 400114 clone MOPC-21), PE mouse IgG2a (Biolegend 400214 clone MOPC-173), and PE mouse IgG2b (Biolegend 400314 clone MPC-11).

### Statistical analysis

All statistical analyses were performed using GraphPad Prism Software (GraphPad Software, La Jolla, CA). A P-value <0.05 was considered significant.

## Results

### Development of an IgG and IgA opsonization assay

The objective of this study was to assess the contribution of NS1-reactive IgG and IgA isotype antibodies to the phagocytic uptake of DENV-infected cells. To this end, we synthesized the previously described DENV-2 reactive murine monoclonal antibody 1G5.3 (30) with either a human IgG1 or an IgA1 Fc domain (**Figure 1A, Supplemental Figure 1**). To confirm that the hIgG1 and hIgA1 isotype conversions did not impact the antigen binding capability of the synthesized mAbs we tested the sNS1 binding activity of the antibodies by ELISA (**Figure 1B**). This analysis showed that the conversion of 1G5.3 to either a human IgG1 or IgA1 isotype did not negatively impact their antigen binding ability, with both mAbs exhibiting an EC_50_ of ~7ng/mL.

**Figure 1:**
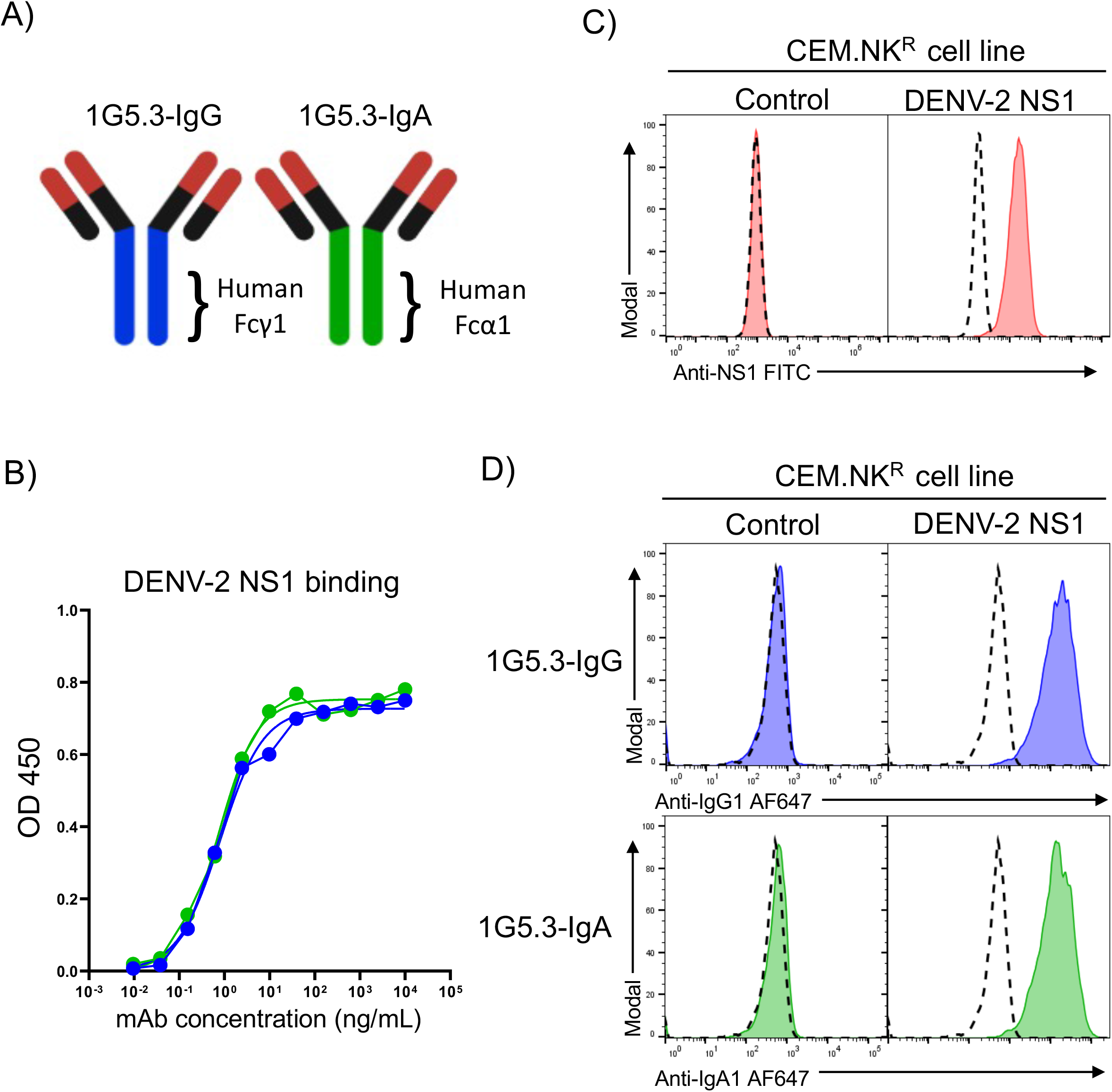
Generation and characterization of a NS1 dependent IgG and IgA opsonization assay. **A)** Schematic representation of the 1G5.3-IgG and 1G5.3-IgA mAbs utilized in this study, **B)** Assessment of sNS1 binding activity of 1G5.3-IgG and 1G5.3-IgA by ELISA. **C)** Characterization of a DENV-2 mNS1-expressing CEM.NK^R^ cell line by staining with the pan-DENV NS1 reactive antibody clone FE8. Dashed histograms indicate staining without primary antibody of the indicated cell type. **D)** Opsonization of DENV-2 NS1-expressing CEM.NK^R^ cells 1G5.3-IgG and 1G5.3-IgA. Dashed histograms indicate staining without primary antibody on the indicated cell line.

To test the ability of 1G5.3-IgG and 1G5.3-IgA to both bind mNS1 and opsonize mNS1 expressing cells we synthesized a DENV-2 NS1 expression constructed that was stably expressed in the CEM.NK^R^ human T lymphoblastoid cell line (**Figure 1C, Supplemental Figure 2**). We confirmed that both 1G5.3-IgG and 1G5.3-IgA exhibited opsonizing activity against the DENV-2 mNS1 expressing CEM.NK^R^ cell line, while exhibiting no binding to the parental line (**Figure 1D**).

### Primary human monocytes are capable of IgG- and IgA-mediated trogocytosis of DENV mNS1-expressing target cells

Having developed both the antibodies and cell lines required to test the ability of IgG and IgA isotype antibodies to facilitate the phagocytic clearance of DENV-infected cells, we next optimized a flow-based antibody-dependent phagocytosis/trogocytosis assay. Antibody-dependent trogocytosis is a process wherein a phagocyte engulfs and degrades a portion of the plasma-membrane of an opsonized cellular target in an FcR dependent fashion. This process – a form of receptor-mediated phagocytosis – allows for professional phagocytes to compromise the membrane integrity of cellular targets larger than themselves, thereby lysing the target cell without having to express and secrete canonical cytolytic factors such as granzyme B and perforin (32–34).

In this assay, target cells were first labeled with the lipophilic dye PKH26 prior to opsonization with either serum or recombinant mAbs (**Figure 2A**). After opsonization, target cells were co-incubated with freshly isolated PMBC from normal healthy donors for 3 hours, after which the phagocytic accumulation of PKH26 labeled target cell membrane within CD14^+^ monocytes was assessed by flow cytometry. In this assay, PKH26 preferentially accumulates in discrete intracellular puncta within CD14^+^ monocytes, consistent with phagocytic uptake (**Supplemental Figure 3**).

**Figure 2:**
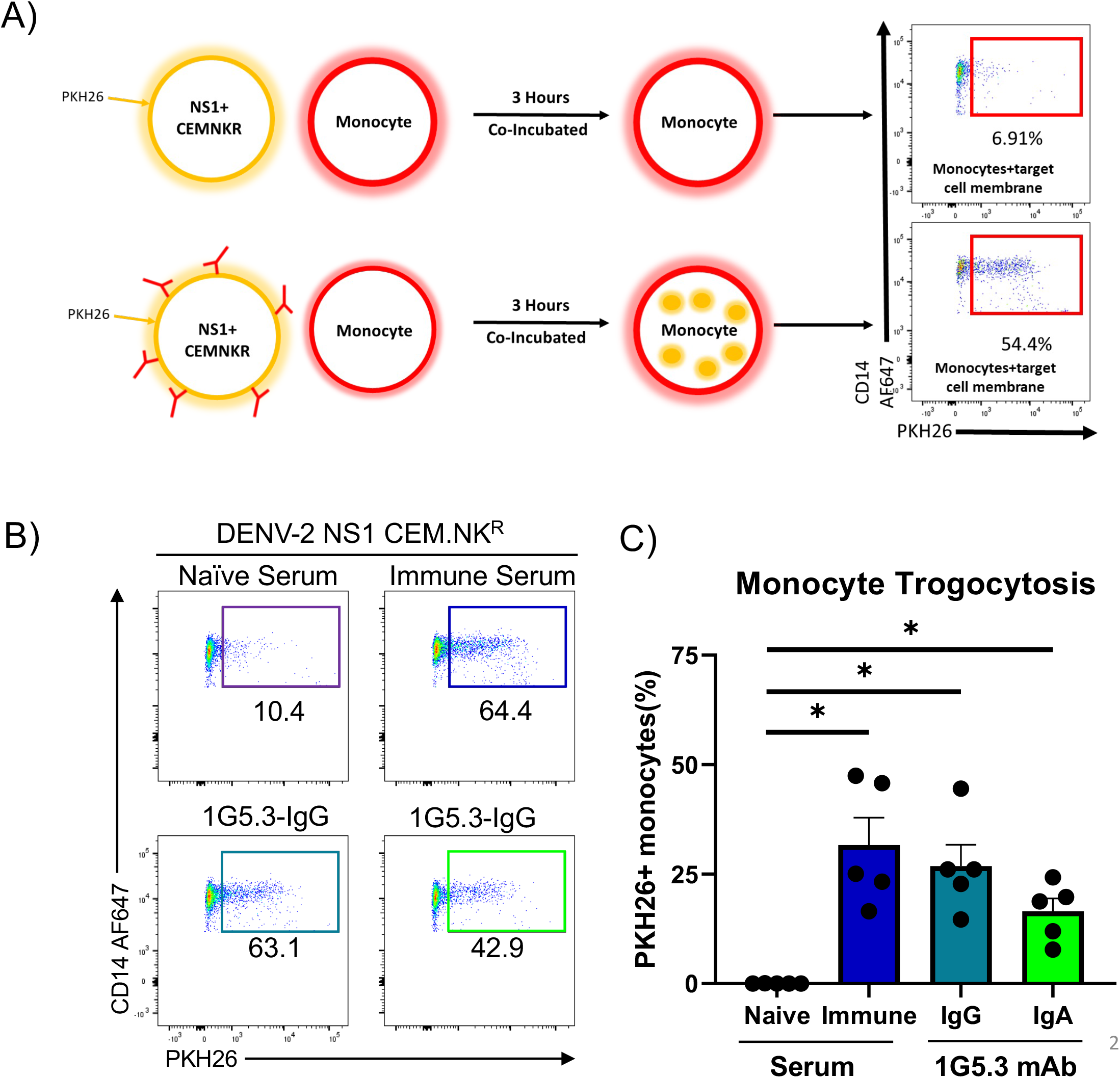
Generation and characterization of a NS1 dependent antibody phagocytosis assay. **A)** Model depicting the antibody-dependent trogocytosis assay used throughout this study, without (top) and with (bottom) opsonizing antibody, **B)** Representative flow plots from the antibody-dependent trogocytosis assay **C)** Percentage of monocytes that were positive for the target cell membrane dye, separated by antibody used to opsonize target, n=5 individual experiments. Error bars are mean ± SEM. * p < 0.05, paired one-way ANOVA.

Using this assay, we assessed the ability of DENV-naïve sera, DENV-immune sera, 1G5.3-IgG and 1G5.3-IgA to facilitate monocytic trogocytosis of DENV-2 mNS1 expressing target cells. While minimal monocytic trogocytosis was observed when the target cells were opsonized with DENV naïve serum, a significant increase in phagocytic activity was observed when the mNS1 expressing target cells were opsonized with either commercially available DENV-immune sera or with either 1G5.3-IgG or 1G5.3-IgA (**Figure 2C**). No monocytic trogocytosis activity was observed in assays using the parental CEM.NKR cell line which lacks DENV-2 NS1 expression as the target cell (**Supplemental Figure 4**). These results indicate that both IgG1 and IgA1 isotype antibodies are capable of facilitating monocytic trogocytosis of DENV mNS1 expressing target cells.

### FcγRI (CD64) and FcαR (CD89) mediate antibody dependent trogocytosis of mNS1 expressing cells

Primary human CD14^+^ monocytes express FcγRIIA (CD32), FcγRI (CD64) and FcαR (CD89), with each Fcγ receptors twice as abundant as the single Fcα receptor (**Figure 3A**). To determine the individual contribution of the Fc receptors on antibody-mediated monocytic trogocytosis we utilized a panel of FcR blocking antibodies to inhibit Fc/FcR interactions during a trogocytosis assay. Pre-treatment with an anti-FcγRIIa blocking antibody (clone IV.3) did not reduce IgG1-medated monocytic trogocytosis (**Figure 3B**). However, pre-treatment with anti-FcγRI (clone 10.1) significantly antagonized IgG dependent trogocytic activity. Neither FcγR blocking antibody reduced IgA mediated trogocytosis under the same assay conditions. However, treatment with the FcαR blocking antibody (clone MIP8a) significantly reduced IgA1-medated monocytic trogocytosis without impacting IgG activity (**Figure 3C**). These data indicate that FcγRI (CD64) and FcαR (CD89) are responsible for facilitating IgG and IgA mediated monocytic trogocytosis of mNS1 expressing cellular material, respectively.

**Figure 3:**
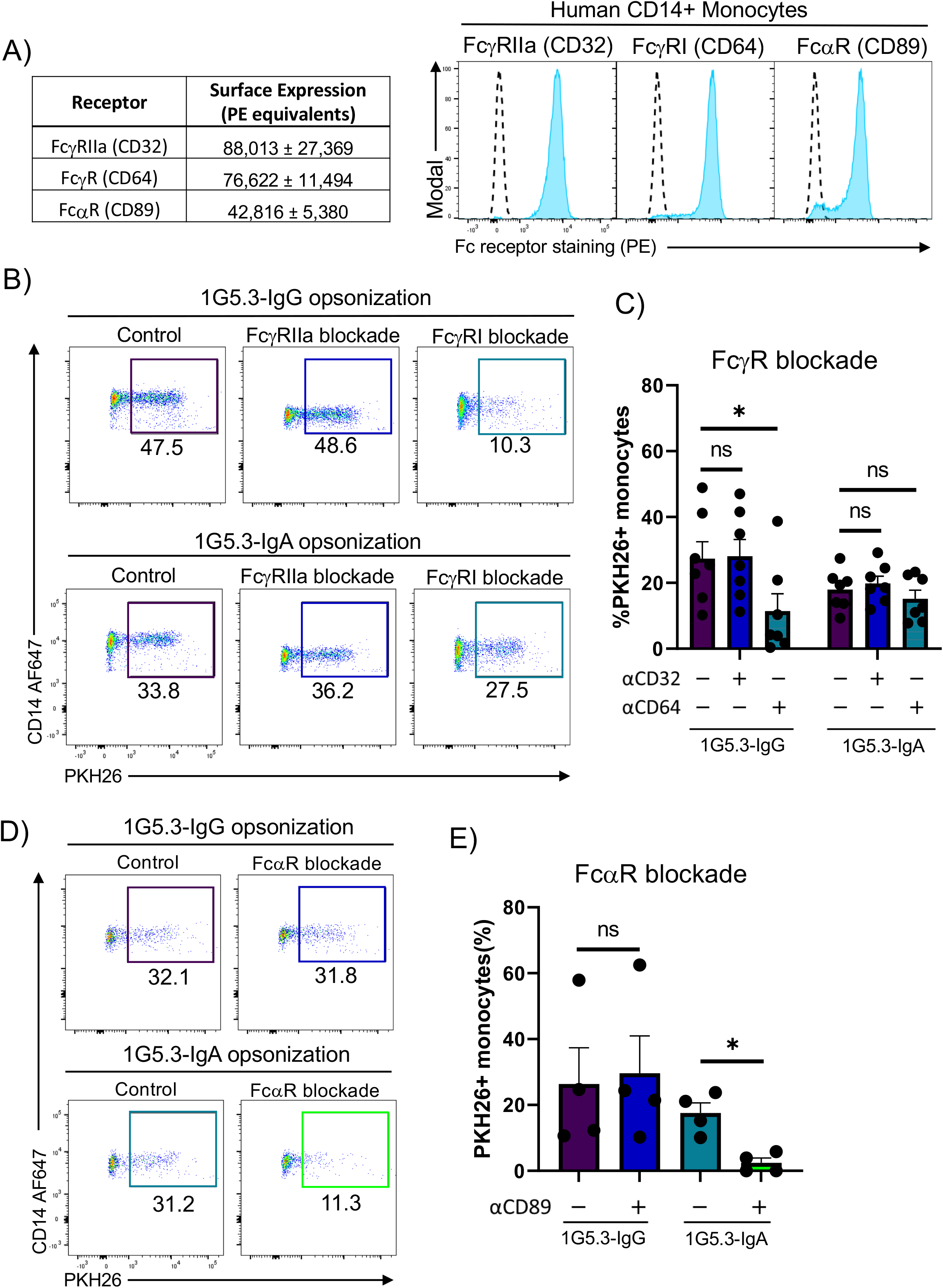
Contribution of FcγRs and FcαR to IgG and IgA mediated phagocytosis. **A)** Surface abundance of Fc receptors on CD14+ monocytes mean ± SEM (left) and representative flow histograms (right). Quantification of Fc receptor abundance was performed utilizing flow cytometric analysis of a standard curve of calibration beads conjugated with known quantifies of PE molecules run in parallel with PBMC stained with saturating concentrations of PE-conjugated anti-FcR antibodies. Dashed line indicates staining of the indicated cell line with a corresponding isotype control. **B)** Representative flow plots of FcγR blocking antibody-dependent trogocytosis assay using both anti-FcγRIIa (CD32) and anti-FcγRI (CD64) antibodies. **C)** Percentage of monocytes positive for target cell membrane indicating antibody-dependent trogocytosis with either anti-FcγRIIa or anti-FcγRI antibodies, n=7 individual experiments **D)** Representative flow plots for FcαR blocking antibody-dependent trogocytosis assay using anti-FcαR antibodies **E)** Percentage of monocytes positive for target cell membrane indicating antibody-dependent trogocytosis with anti-FcαR antibodies, n=4 individual experiments. All error bars are mean ± SEM. * p < 0.05, paired one-way ANOVA

### No evidence of synergy between IgG and IgA antibodies in mediating monocytic trogocytosis

Having established both IgG and IgA isotype antibodies can independently mediate monocytic trogocytosis of mNS1 expressing cells, we next endeavored to asses any potential synergistic activity of NS1-specific IgG and IgA in this process. It has been previously observed that IgA is able to enhance IgG-mediated ADCC by monocytes and PMNs, and we hypothesized a similar effect might be observed in the setting of antibody dependent monocytic trogocytosis (35). To this end we titrated both 1G5.3-IgG and 1G5.3-IgA independently or mixed at a 1:1 ratio to determine if the mix of antibodies exhibited any differential activity when used together. Both 1G5.3-IgG and 1G5.3-IgA facilitated nearly identical levels of trogocytic activity across more than 3 logs of antibody concentration, as did the 1:1 mixture of the two antibodies used at the same opsonizing concentration (**Figure 4**). These results suggest that IgG and IgA can act in an additive fashion to facilitate monocytic trogocytic of antigen-expressing cells, but there is no evidence of synergistic activity.

**Figure 4:**
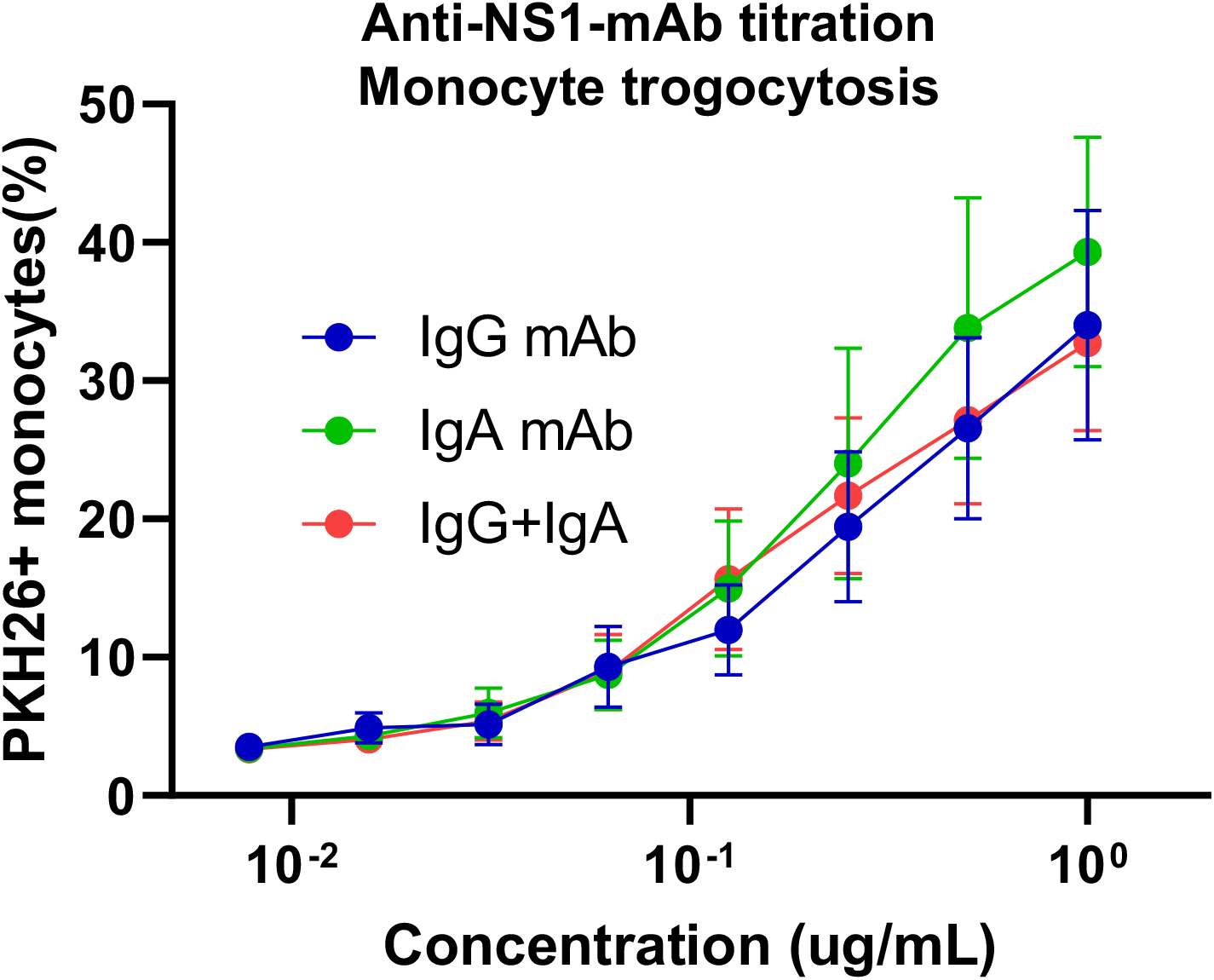
Assessment of IgG and IgA synergism in antibody-mediated phagocytosis. **A)** Titration curve for monocytic trogocytosis using either 1G5.3-IgG (blue), 1G5.3-Iga (green), or a 1:1 mix of 1G5.3-IgG and 1G5.3-IgA at the indicated total antibody concentration. n=3 individual experiments

### IgA and IgG facilitate antibody-dependent monocytic uptake of DENV-2 infected cells

The analysis presented in the study thus far has utilized a transgenic expression system to uniformly express mNS1 on a target cell. While this represents a well-controlled and highly reproducible experimental system, we wanted to confirm that the results obtained thus far were still observed in cells expressing mNS1 as the result of DENV infection rather than the result of enforced transgenic expression of the protein. In a natural infection, not all cells are infected and there is variable mNS1-expression on the infected cells which may influence the efficiency of the process. To better study the ability of monocytes to phagocytose DENV-2 infected cells utilizing NS1-reactive antibodies, we infected DC-SIGN expressing CEM.NK^R^ cells with DENV-2 for use as target cells in our antibody-dependent trogocytosis assay (**Figure 5A**). DC-SIGN expressing CEM.NK^R^ cells were infected with DENV-2 overnight, harvested, then stained with either 1G5.3-IgG or 1G5.3-IgA followed by anti-human IgG and IgA fluorescent antibodies to verify NS1 expression (**Figure 5B**). Approximately 25% of the target cells expressed mNS1 in these conditions at levels consistent with those observed in the DENV-2 NS1 expressing CEM.NK^R^ cell line. Having confirmed mNS1 expression on these DENV-2 infected cells, they were used in the antibody-dependent trogocytosis assay described above in place of the NS1-expressing CEM.NK^R^ cell line. As was observed with the mNS1 expressing CEM.NK^R^ cell line, DENV-immune serum, 1G5.3-IgG and 1G5.3-IgA were observed to facilitated monocytic trogocytosis of DENV-2 infected cells (**Figure 5C**).

**Figure 5:**
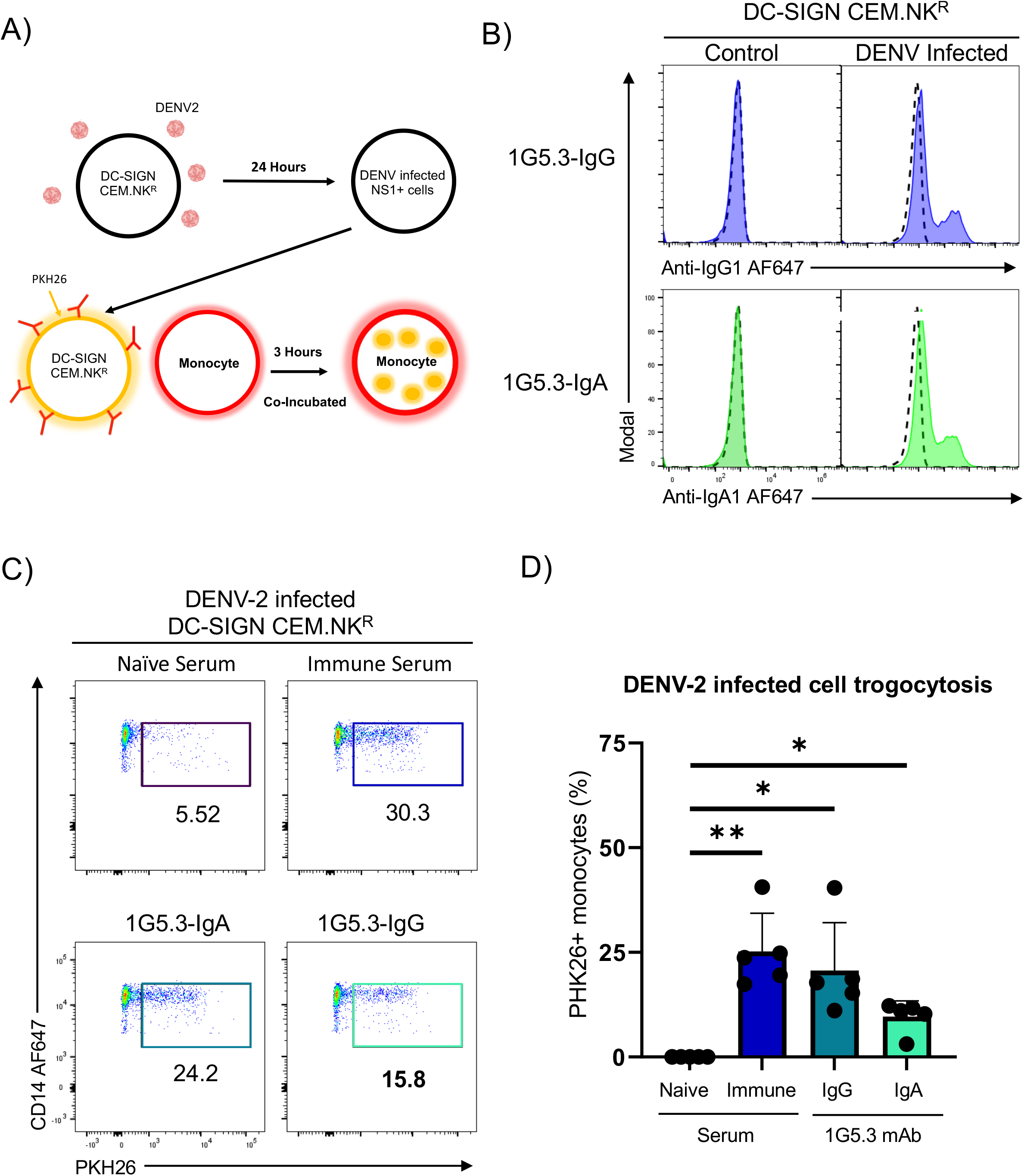
IgA and IgG facilitate antibody-dependent monocytic uptake of DENV-2 infected cells. Model depicting the antibody-dependent trogocytosis assay including the infection of DC-SIGN expressing CEM.NK^R^ cells, **B)** Opsonization of DENV-2 infected DC-SIGN expressing CEM.NK^R^ cells with NS1-reactive 1G5.3-IgG and 1G5.3-IgA mAbs, with percent monocytes positive for target cell membrane indicating antibody-dependent trogocytosis. Dashed line in histograms indicates staining with DENV naïve serum. **C)** Representative flow plot for antibody-dependent trogocytosis assay using DENV-2 infected DC-SIGN expressing cells as targets **D)** Percentage of monocytes positive for target cell membrane indicating antibody-dependent trogocytosis with DENV-2 infected cells as targets, n=5 individual experiments. Error bars are mean ± SEM. * p < 0.05, ** p < 0.01, paired one-way ANOVA

### Soluble NS1 is capable of abrogating monocyte antibody-dependent trogocytosis

While our study thus far has focused on the immunologic contribution of mNS1 in dengue, NS1 is best known for its sNS1 form which can be found at microgram/ml quantities in serum during acute DENV infection (36–38). It is currently unclear why flaviviruses have evolved to secrete such high levels of NS1 into the extracellular space. While sNS1 has been described as a potential pathogenic factor capable of triggering multiple immunopathogenic programs, there is no clear advantage to the virus from an evolutionary standpoint. However, it has been proposed that sNS1 acts to aid in immune avoidance by depleting complement in the vicinity of DENV-infected cells (39). In light of the results obtained in our study this far, we hypothesized that another reason why DENV-infected cells may produce such abundant sNS1 was to act as immunologic “chaff”, protecting DENV-infected cells from opsonizing and subsequent clearance by NK cells and phagocytes.

To test this hypothesis, we modified our monocytic antibody-dependent trogocytosis assay to include the addition of purified recombinant NS1 during the opsonization and co-incubation steps. The CEM.NK^R^ cell line used in this study produces very little sNS1 following either transfection with the NS1 expression construct or following infection with DENV, so exogenous recombinant sNS1 was spiked into the cell cultures to better mirror the conditions observed during natural DENV infection. Given the highly preferential DENV-2 NS1 binding activity of the 1G5.3 mAbs used in this study, DENV-1 NS1 was used as a control to account for any immunomodulatory effects of NS1 itself outside of its role as an antibody target. Using this modified assay, we observed that the addition of purified DENV-2 NS1 was able to abrogate the monocytic antibody-dependent trogocytosis of DENV-2 NS1-expressing CEM.NK^R^ cells (**Figure 6A**). This process appears was specific to the anti-NS1 antibodies because the addition of serotype mismatched DENV-1 NS1 had no effect on the level of antibody-dependent trogocytosis (**Figure 6A**).

**Figure 6:**
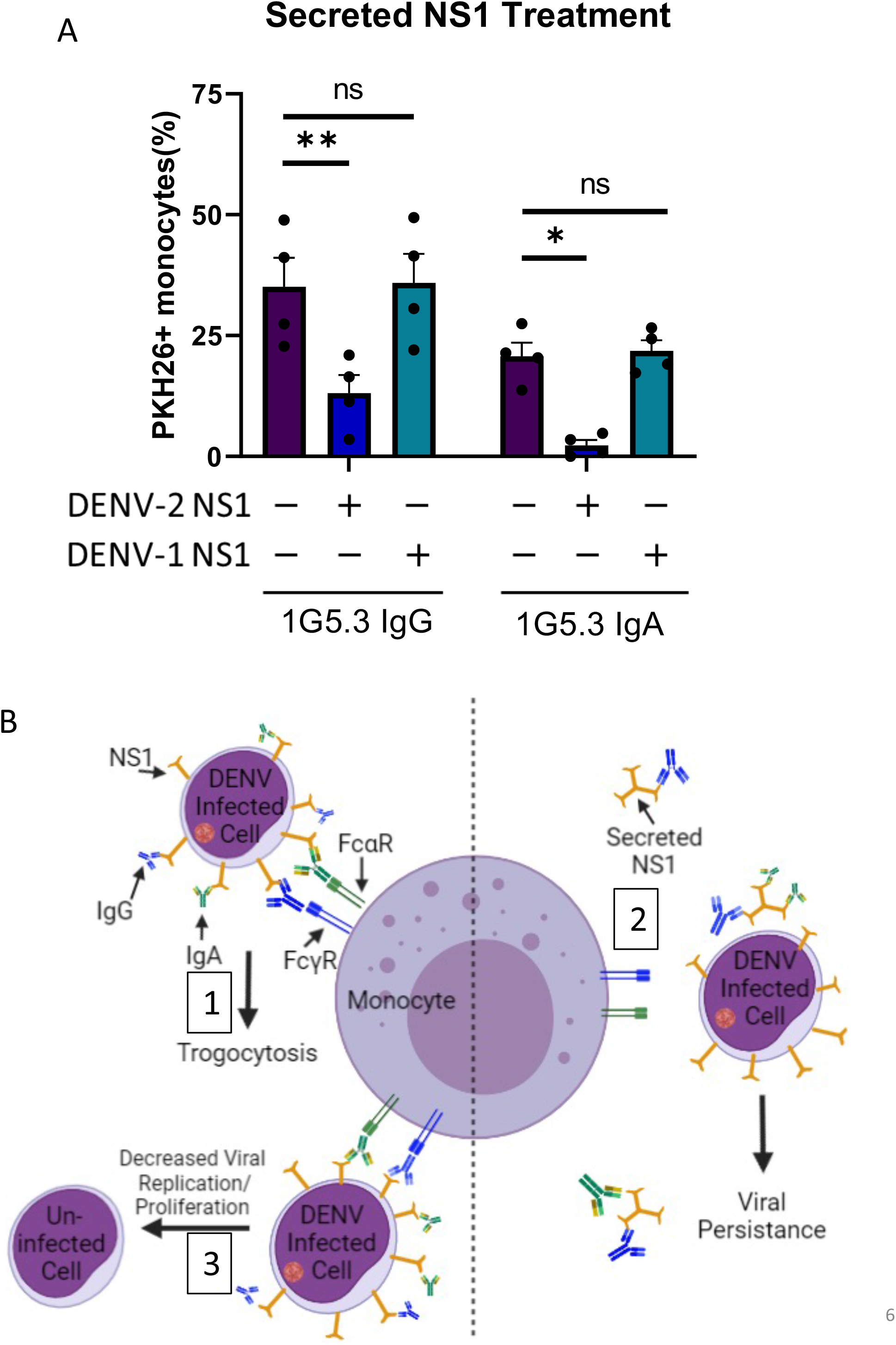
Soluble NS1 is capable of antagonizing monocytic antibody-dependent trogocytosis. **A)** Trogocytosis assay with purified secreted DENV-1 or DENV-2 NS1 added during the opsonization step. Percent monocytes positive for target cell membrane indicates trogocytosis, n=4 individual experiments. Error bars are mean ± SEM. * p < 0.05, ** p < 0.01, paired one-way ANOVA **B)** Proposed model for the role of non-neutralizing antibodies in clearing DENV-infected cells.

The model shown in **Figure 6b** depicts the proposed mechanism for functional role of NS1-reactive antibodies. NS1 expressed on the cell surface of infected cells and is opsonized by NS1-reactive IgG and IgA antibodies (**Fig. 6b, step 1**). However, this trogocytosis can be abrogated by secreted NS1 binding NS1-reactive antibodies, preventing opsonization (**Fig. 6b, step 2**). We propose that these NS1-reactive antibodies mediating antibody-dependent trogocytosis lead to decreased viral replication; however, investigation is needed (**Fig 6b, step 3**).

## Discussion

In this study, we defined a role for NS1-reactive IgG and IgA antibodies in mediating antibody-dependent monocytic trogocytosis of DENV mNS1 expressing cells. Our analysis revealed that both IgG and IgA isotype antibodies are capable of facilitating phagocytic uptake of DENV infected material, and that these two antibody isotypes can work together in an additive – but not synergistic – fashion when combined. Notably, we observed that the addition of sNS1 dramatically antagonized the ability of both NS1-reactive IgG and IgA to facilitate monocytic uptake of NS1 expressing cellular material, highlighting a potential mechanism of DENV immune evasion.

While our study focuses on DENV, NS1-reactive antibodies have been observed in circulation after infection with numerus other flavivirus including Zika virus (ZIKV) (40–43). Notably, ZIKV NS1-reactive antibodies have been shown to be capable of facilitating NK cell degranulation and lysis of NS1 expressing cells using a system similar to that presented in our study (40). NS1 reactive antibody therapies and NS1 containing vaccine products have all exhibited promise in preclinical testing, with these products exhibiting the ability to facilitate both ADCC as well as complement fixation on flavivirus infected cells (44–47). These observations highlight the diversity Fc dependent effector mechanism by with non-neutralizing/NS1-reactive antibodies may help resolve or prevent acute flavivirus infections, but are in no way exhaustive. Finally, in addition to facilitating a broad array of effector function, NS1 reactive antibodies have the additional benefit of having no potential infection enhancement activity, thereby circumventing many of the historic risks associated with vaccines and/or monoclonal antibodies that target structural proteins of the flavivirus virion.

We proposed that sNS1 plays a role in DENV immune evasion by sequestering anti-NS1 antibodies to prevent opsonization of DENV-infected cells, thereby shielding them from clearance via monocytic antibody-dependent trogocytosis, ADCC, or other Fc receptor mediated effector mechanisms. There are many examples of secreted viral proteins playing an immune avoidance role for the virus. Cells infected with Cowpox Virus secrete the CPXV14 protein that prevents antibody-mediated T-cell activation by binding to FcγR (48).Vaccinia virus also encodes a secretory protein, VCP, that binds C3b and C4b to inhibit complement activation (49). More closely related to DENV, West Nile virus NS1 has been shown to inhibit complement activation by binding to regulatory protein factor H (50). This is similar to how DENV sNS1 depletes complement near DENV-infected cells (39, 51). DENV NS1 has been shown to either independently, or as a complex with vitronectin, block the formation of the membrane attack complex (51). DENV, West Nile, and Zika virus NS1 are able to block C9 polymerization (51). Understanding other potential immune avoidance roles DENV sNS1 plays and the mechanisms behind them could help to better treat DENV-infected patients, possibly using antibodies against NS1 as a therapeutic intervention.

There are a few limitations that should be considered while interpreting the results of this study. Most of the data were generated using a cell line that was modified to stably express NS1 on the cell surface instead of infected cells. The consequence of this is that every target cell was mimicking an infected cell whereas in a natural infection most cells are not infected. This also exaggerated the percentage of monocytes that had internalized target cell membrane. The use of monoclonal antibodies specific to NS1 presents the problem of not using physiological concentrations of NS1-reactive antibody. The use of polyclonal DENV-immune plasma from DENV-infected patients showed that the monoclonal antibodies mediated similar levels of trogocytosis by monocytes, but additional work is required to determine the impact of serum-level concentration of anti-NS1 in a more complex immune setting.

In closing, we suggest that our study highlights several unappreciated facets of anti-DENV immunity. Not only have we demonstrated a role for NS1-reactive IgA in the setting of acute flavivirus infection, but have identified a putative method of flavivirus immune evasion through sNS1. These results will inform not only future countermeasure development, but may also represent a new correlate of risk or protection from infection.

## ACKNOWLEDGEMENTS

We gratefully acknowledge technical assistance provided by Lisa Phelps of the SUNY Upstate Medical University Flow Cytometry Core, to Dr. Joel Wilmore for his critical review of the manuscript, and to Dr. Nathan Roy for his technical assistance with microscopy. The opinions or assertions contained herein are the private views of the authors and are not to be construed as reflecting the official views of the US Army or the US Department of Defense. Material has been reviewed by the Walter Reed Army Institute of Research. There is no objection to its presentation and/or publication. The investigators have adhered to the policies for protection of human participants as prescribed in AR 70-25.

## Author contributions. Conceptualization

M.J.W., J.R.C., A.T.W., **Formal analysis:** M.J.W., A.T.W **Funding acquisition:** A.T.W. **Investigation:** M.J.W., A.D.W, L.E.B. **Resources:** J.R.C., A.T.W. **Writing – original draft:** M.J.W., A.T.W **Writing – review & editing:** all authors

