## Supplemental figures for "Soluble NS1 antagonizes IgG- and IgA-mediated monocytic phagocytosis of DENV infected cells"

A

1G5.3 Light Chain:

MGWSCIILFLVATATGVHSEIVLTQSPASLAVSLGQRATISCRASESVEYSGTSLMHWYQQK
PGQPPKLLIYAASNVESGVPARFSGSGSGTDFSLNIHPVEEDDIAMYFCQQSRKVPYTFGGG
TKLELKRTVAAPSVFIFPPSDEQLKSGTASVVCLLNNFYPREAKVQWKVDNALQSGNSQESV
TEQDSKDSTYSLSSTLTLSKADYEKHKVYACEVTHQGLSSPVTKSFNRGEC.

B 1G5.3 IgG Heavy Chain:

MGWSCIILFLVATATGVHSEVKLQESGPGLVRPSQSLSLTCSVTGYSITSGYYWNWIRQFPG
NKLEWMGYISYDGRSNYNPSLKNRISITRDTSKNQFFLKLNFVTTEDTATYYCASFYYYTSR
PLVYWGQGTLLTVSSASTKGPSVFPLAPSSKSTSGGTAALGCLVKDYFPEPVTVSWNSGALT
SGVHTFPAVLQSSGLYSLSSVVTVPSSSLGTQTYICNVNHKPSNTKVDKKVEPKSCDKTHTC
PPCPAPELLGGPSVFLFPPKPKDTLMISRTPEVTCVVVDVSHEDPEVKFNWYVDGVEVHNAK
TKPREEQYNSTYRVVSVLTVLHQDWLNGKEYKCKVSNKALPAPIEKTISKAKGQPREPQVYT
LPPSRDELTKNQVSLTCLVKGFYPSDIAVEWESNGQPENNYKTTPPVLDSDGSFFLYSKLTV
DKSRWQQGNVFSCSVMHEALHNHYTQKSLSLSPGK

C 1G5.3 IgA Heavy Chain:

MGWSCIILFLVATATGVHSEVKLQESGPGLVRPSQSLSLTCSVTGYSITSGYYWNWIRQFPG
NKLEWMGYISYDGRSNYNPSLKNRISITRDTSKNQFFLKLNFVTTEDTATYYCASFYYYTSR
PLVYWGQGTLLTVSSASPTSPKVFPLSLCSTQPDGNVVIACLVQGFFPQEPLSVTWSESGQG
VTARNFPPSQDASGDLYTTSSQLTLPATQCLAGKSVTCHVKHYTNPSQDVTVPCPVPSTPPT
PSPSTPPTPSPSCCHPRLSLHRPALEDLLLGSEANLTCTLTGLRDASGVTFTWTPSSGKSAV
QGPPERDLCGCYSVSSVLPGCAEPWNHGKTFTCTAAYPESKTPLTATLSKSGNTFRPEVHLL
PPPSEELALNELVTLTCLARGFSPKDVLVRWLQGSQELPREKYLTWASRQEPSQGTTTFAVT
SILRVAAEDWKKGDTFSCMVGHEALPLAFTQKTIDRLAGKPTHVNVSVVMAEVDGTCY

**Supplemental Figure 1: Sequences of NS1 reactive monoclonal antibodies.** NS1-Reactive monoclonal antibody sequences for the light chain (A), heavy chain for IgG (B), and heavy chain for IgA (C)

### DENV-2 NS1-NS2A sequence

```
MNSRSTSLSVTLVLVGIVTLYLGVMVQADSGCVVSWKNKELKCGSGIFITDNVHTWTEQYK
FQPESPSKLASAIQKAHEEGICGIRSVTRLENLMWKQITPELNHILSENEVKLTIMTGDIK
GIMQAGKRSLRPQPTELKYSWKTWGKAKMLSTESHNQTFLIDGPETAECPNTNRAWNSLEV
EDYGFGVFTTNIWLKLKEKQDVFCDSKLMSAAIKDNRAVHADMGYWIESALNDTWKIEKAS
FIEVKNCHWPKSHTLWSNGVLESEMIIPKNLAGPVSQHNYRPGYHTQITGPWHLGKLEMDF
DFCDGTTVVVTEDCGNRGPSLRTTTASGKLITEWCCRSCTLPPLRYRGEDGCWYGMEIRPL
KEKEENLVNSLVTAGHGQVDNFSLGVLGMALFLEEMLRTRVGTKHAILLVAVSFVTLITGN
MSFRDLGRVMVMVGATMTDDIGMGVTYLALLAAFKVRPTFAAGLLLRKLTSKELMMTTIGI
VLLSQSTIPETILELTDALALGMMVLKMVRNMEKYQLAVTIMAILCVPNAVILQNAWKVSC
TILAVVSVSPLLLTSSQQKTDWIPLALTIKGLNPTAIFLTTLSRTSKKR
```

**Supplemental Figure 2:** DENV-2 sequence used in the NS1-expressing CEM.NK$^R$ cell line. Yellow = signal peptide, green = NS1, blue = NS2A

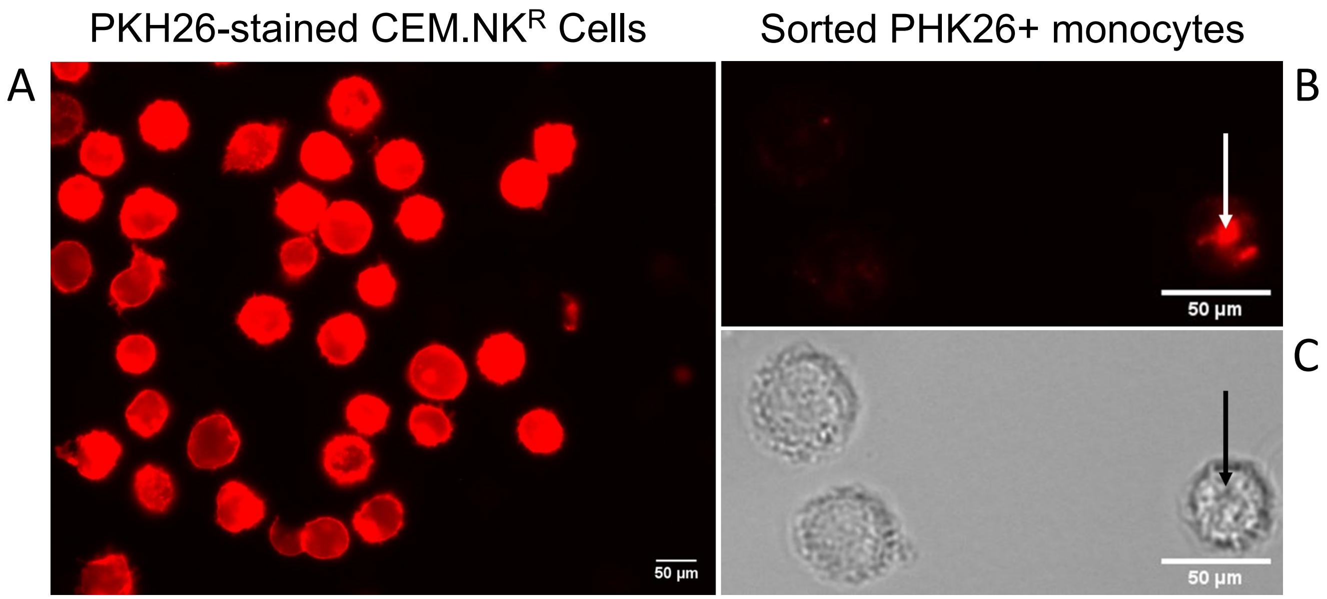


**Supplemental Figure 3:** Micrographs of PKH26 stained CEM.NKR cells and sorted PHK26+ CD14+ monocytes

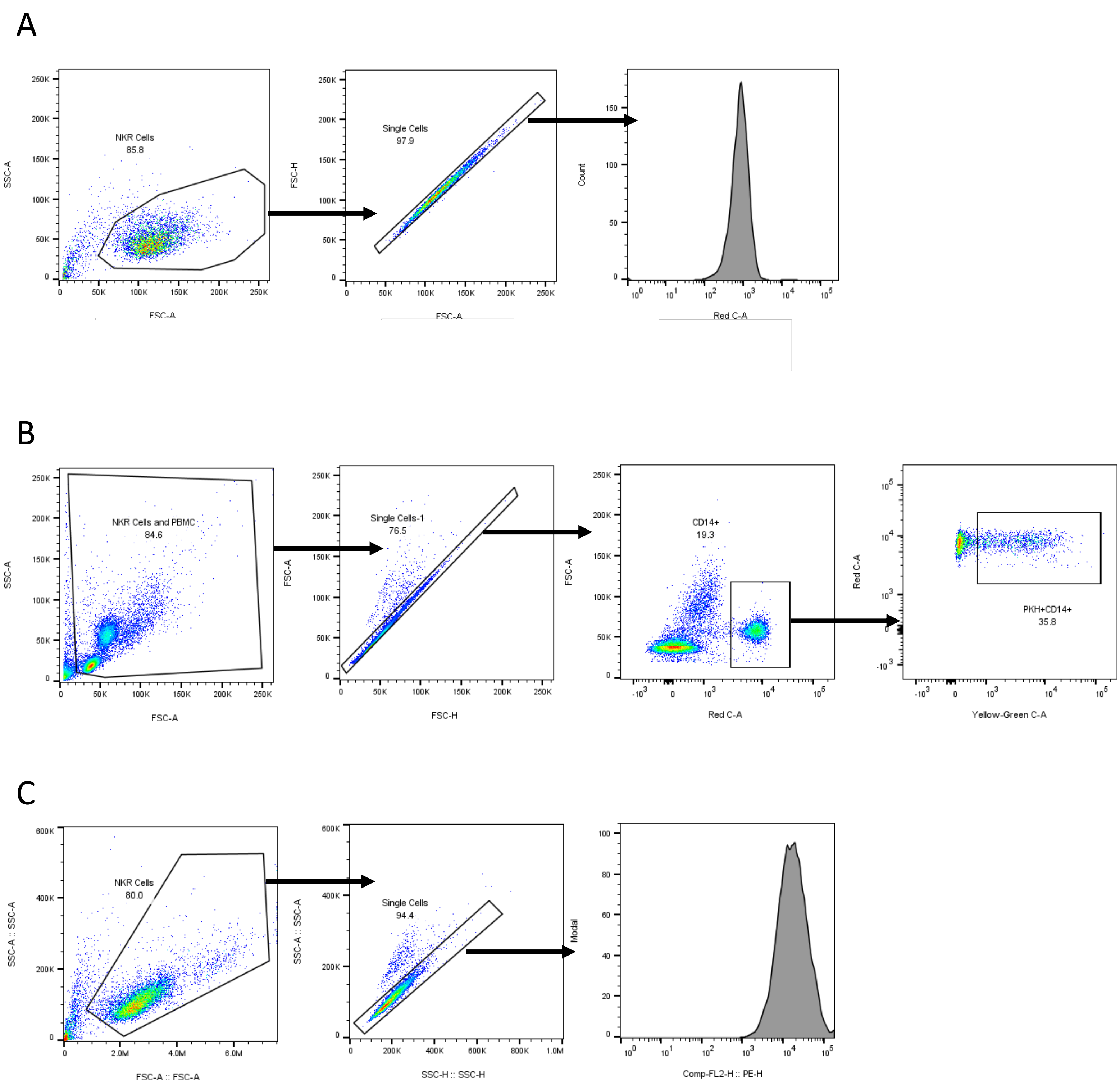


**Supplemental Figure 4:** A) Gating layout used for opsonization assays of NS1-expressing CEM.NK$^R$ cells B) Gating layout used for antibody-dependent trogocytosis assays, C) Gating layout used for opsonization of DC-SIGN expressing CEM.NK$^R$ cells

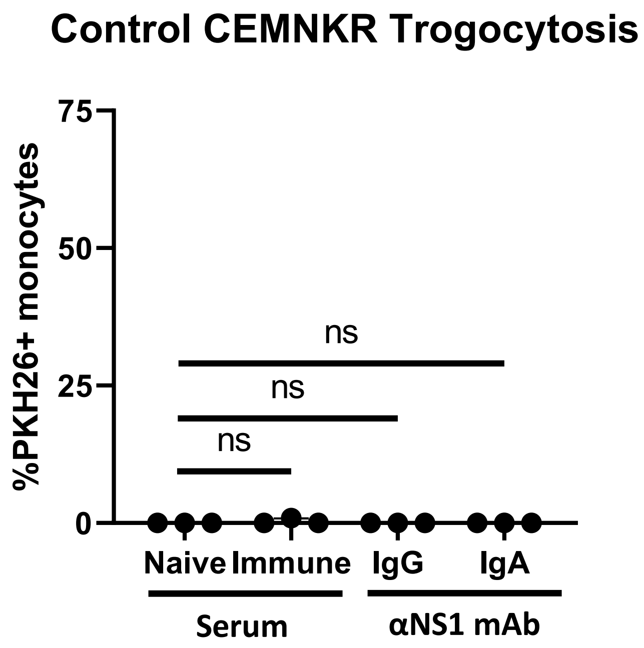


**Supplemental Figure 5:** Trogocytosis assay results using the parental CEM.NK$^{R}$ cells as targets. n=3 individual experiments

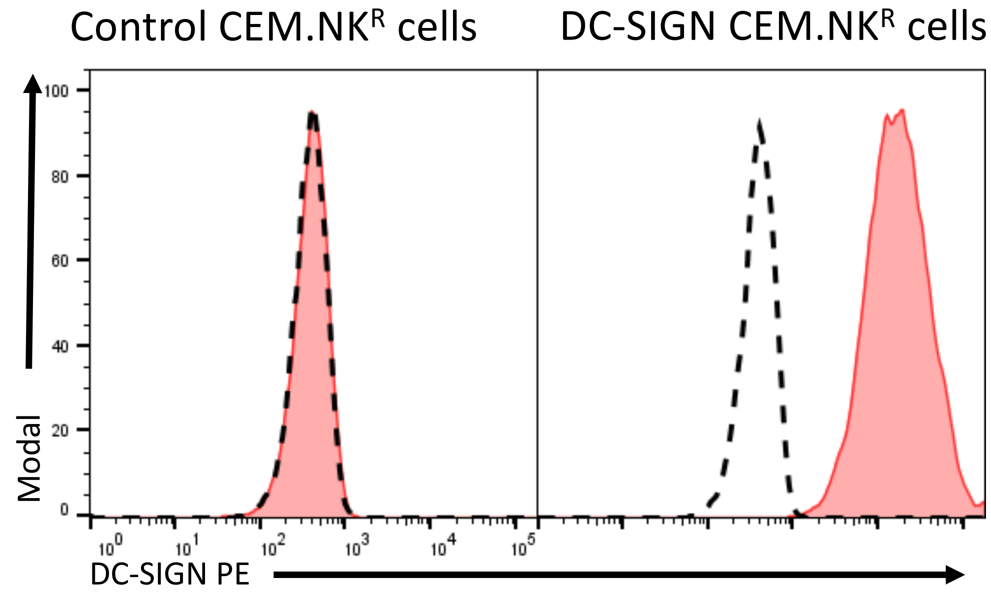


**Supplemental Figure 6:** Surface DC-SIGN expression on control and DC-SIGN expressing CEM.NK$^R$ cells. Dashed line indicates unstained cells.

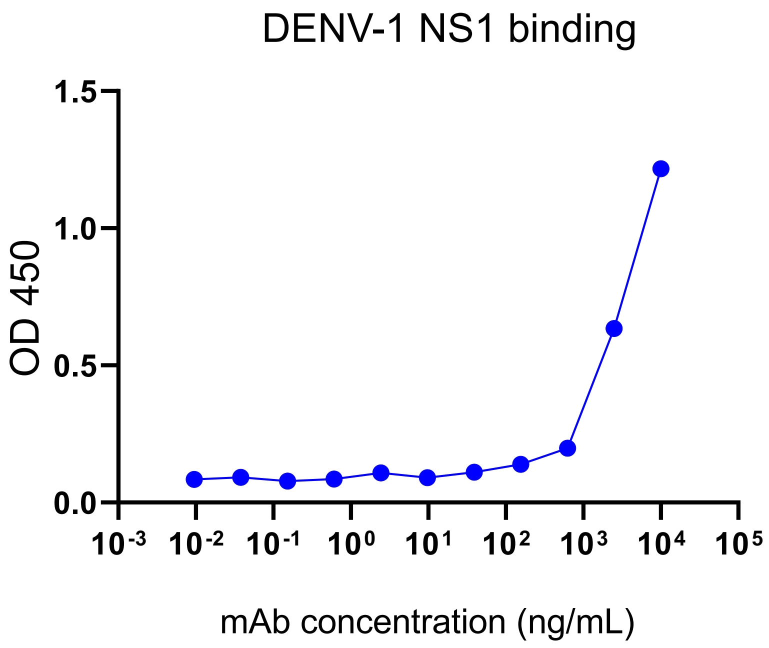


**Supplemental Figure 7:** DENV-1 NS1 binding of 1G5.3 IgG as assessed by ELISA
